# Disentangling the effects of past logging and ongoing cryptic anthropogenic disturbance on vegetation structure and composition in Himalayan Foothills, India

**DOI:** 10.1101/2021.04.21.440725

**Authors:** Monica Kaushik, Sutirtha Dutta, Gopal S. Rawat, Pratap Singh, Dhananjai Mohan

**Affiliations:** Wildlife Institute of India, Chandrabani, Dehradun-248001, India; Uttarakhand Forest Department

**Keywords:** Chronic-anthropogenic disturbance, *Lantana camara*, Regeneration failure, Selective-logging

## Abstract

Most tropical forests have undergone commercial logging. Even where logging has ceased, subsistence harvest of forest resources often persists especially in South-East Asia. Understanding of impacts of frequent forest resources extraction in areas recovering from past selective logging would be essential for designing the appropriate management interventions.

We studied the impacts of current chronic anthropogenic disturbances (hereafter CAD) and past selective logging on vegetation structure, diversity, and regeneration, and the invasion of a non-native shrub, *Lantana camara*, in three major forest types in the Himalayan foothills, India. We analyzed field data on intensity of CAD and vegetation variables, collected from 269 stratified random plots, using ordination and generalized linear (mixed) modeling approaches.

Our results, based on 2758 adult trees of 54 species, showed that forest types differed in disturbance regimes depending on protection level and availability of fodder tree species. Intensity of CAD depended on proximity to settlements (for livestock related disturbances). Whereas selective logging, including firewood collection, was associated with land protection status. Selective logging reduced the extent of mature forest but facilitated regeneration, thereby promoting secondary forest features such as tree density and canopy cover. In contrast, the interaction between lopping and selective logging was negatively associated with regeneration. Past logging facilitated *L. camara* invasion in Dry and Hill forests but not in Moist forest. Finally, while selective logging marginally enhanced tree diversity, CAD reduced native shrub diversity.

Our study demonstrates that selective logging followed by CAD arrest forest recovery, as evident from the suppression of mature forest elements, loss of shrub biomass, reduced regeneration rate, and facilitation of invasive species. To abate these impacts, alternative livelihood/subsistence options that sustain forests and local communities should be explored. Additionally, CAD management should be site-specific as local ecological contexts modify their impacts on forests.

## 1. Introduction

Tropical forests have been historically exploited through selective logging at large-scales by the timber industry and at small scales by forest dwelling and adjoining rural communities for their sustenance and livelihoods (Geist and Lambin, 2002). The most easily accessible forests have undergone commercial selective logging in the past, leading to changes in wildlife habitat and ecological successions at local scales. Selective logging is now illegal in many countries and has largely ceased within Protected Areas (Bertzky et al., 2012). However, small-scale illegal selective logging still persist in some developing countries to meet local demands (Chazdon, 2014; Yonariza and Webb, 2017). Furthermore, extraction of forest resources (i.e. collection of fuel wood, fodder, and non-wood forest products such as medicinal plants) by local communities occur almost daily (Davidar et al., 2010; Singh, 1998).

Chronic anthropogenic disturbances (CADs) in the form of small-scale and continuous extraction of forest resources is prevalent in many tropical forests, particularly in developing countries (Singh, 1998). Such disturbances can substantially modify habitat structure, alter species composition and threaten the persistence of local flora and fauna (Ribeiro-Neto et al., 2016; Shahabuddin and Kumar, 2006). Impacts on faunal communities can trickle down to alter regulatory ecosystem services such as seed dispersal and pollination (Leal et al., 2014).

Loss or thinning of tree canopy due to lopping and removal of fuel wood can modify microclimatic conditions such as ambient temperature, relative humidity, light and movement of wind (Sekercioglu, 2002) that in turn can affect flower production, pollination, seed dispersal and regeneration of native trees and shrubs. Opening of the tree canopy can facilitate the establishment of pioneer species or light demanding invasive alien plants (IAPs), suppressing native species and homogenizing local biodiversity (Gordon, 1998). Yet, quantification of the effects of such large and small-scale disturbances on vegetation structure and composition has typically been restricted to one or two types of forests (Ramírez-Marcial et al., 2001; Shahabuddin and Kumar, 2006). Previous research have strongly advocated conducting exhaustive analysis of such drivers across various forest types at multiple scales (Saberwal and Rangarajan, 2005; Shahabuddin and Prasad, 2004).

Tropical and sub-tropical forests along the Himalayan foothills are of high conservation significance owing to diverse forest types, rich biodiversity and higher carbon sequestration potential (Sharma et al., 2010; Sivakumar et al., 2010). These forests support populations of conservation flagships such as the tiger *Panthera tigris* and Asian elephant *Elephas maximus*, and meet subsistence requirements of a large number of local communities and their domestic livestock (Robinson et al., 2014). The forests along the western Himalayan foothills have been subjected to CAD from the local agro-pastoral as well as migratory pastoral communities (e.g., *Gujjars, Gaddis* and *Bakkarwals*). These communities have been utilizing the foothill forests for fodder, fuelwood and construction material for at least 110 years (Gooch, 2009). Some parts of these forests were brought under the Protected Area network by state Governments to conserve native biodiversity and wildlife. Other areas are partially protected as Reserve Forests, which were used for selective logging of timber and subsequent plantation of commercially important trees such as *Tectona grandis, Eucalyptus sp*., *Bombax ceiba*. and *Ailanthus excelsa*. Collectively these forests represent a mosaic of protected lands and areas subject to heavy forest resource extraction by local communities, resulting in a gradient of anthropogenic pressures, and providing a unique opportunity to study their impacts on forest ecology.

Here, we evaluate the ecological consequences of such disturbances across three dominant forest types within the Shivalik ranges of the Himalayan foothills. Specifically, we addressed the following questions: 1) How does intensity of anthropogenic pressures vary across major forest types and management regimes? 2) How do past selective logging and current CADs, alone or in combination, affect vegetation structure, floral diversity, and forest age structure across forest types? 3) Are these disturbances associated with the invasion of the exotic shrub, *Lantana camara* in the study area*?*

## 2. Methods

### 2.1 Study Area

We conducted this study in the forests of Himalayan foothills (29°57’ to 31°20’ N and 77°35’ to 79°20’E). Study region is 74 km long and 14 km wide and includes flat alluvial plains and highly dissected Shivalik ranges between two major rivers, Yamuna in the west and Ganga in the east (Fig. 1). We restricted our sampling to this region to control for climatic, physical and edaphic features that might cause variation in vegetation characteristics. The Shivalik ranges run parallel to the lesser (outer) Himalaya in both northwestern and southeastern directions, which results in varying slopes and aspects across the region. The study area lies in the subtropical zone and has a moderate monsoon climate. Annual rainfall ranges between 1600-1800 mm, most of which falls during July-September from the southwest monsoon. Altitude of the study area varies from 300-1500 m. The vegetation falls under the broad category of Tropical Moist Deciduous forests (Champion and Seth, 1968), which are dominated by Sal tree (*Shorea robusta*), but also includes 11 finer forest types: Moist Shivalik Sal, Dry Shivalik Sal, Dry plain Sal, Mixed Deciduous, Mixed Scrub, Himalayan moist scrub, Subtropical Pine, Low Alluvial Savannah, Khair-sissoo, Riverine and Hill Valley Swamp, and Plantations.

**Figure 1:**
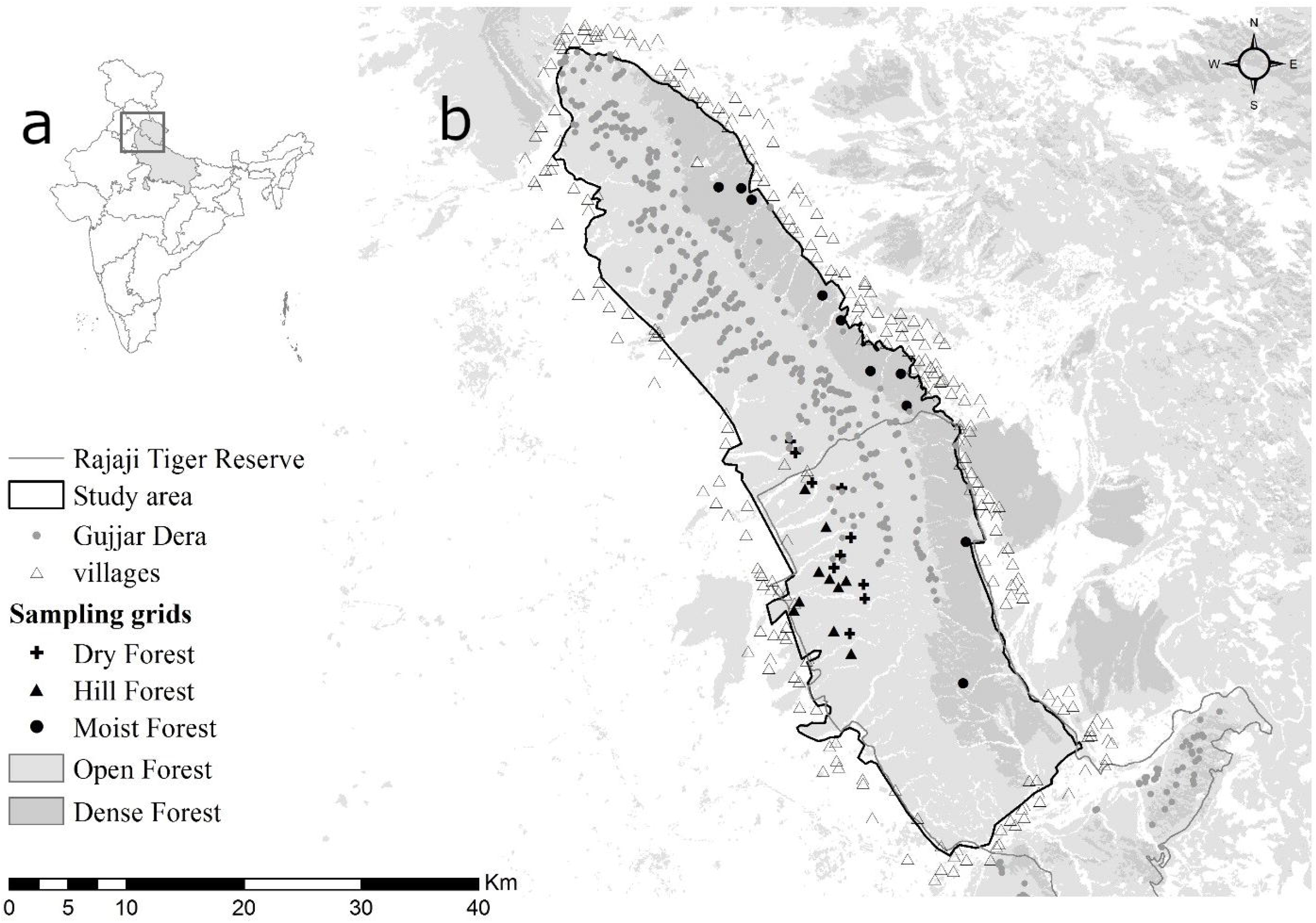
(a) Location of the study area within two Indian states in the Himalayan Foothill region. (b) Location of the 30 study sites where the vegetation community was assessed within three distinct forest types (Dry, Hill and Moist forest). Human presence in the study area is evident in form of small hamlets (filled gray circles) and villages (open triangles).

We selected the following three forest types for sampling due to their maximum spatial coverage in this landscape:

- Dry plain Sal Forest (hereafter Dry Forest): Forests located on drier, gentle slopes and flat areas on the southern slopes of Shivalik dominated by *S. robusta* and associated plants (*Terminalia tomentosa, T. belerica* and *Lagerstroemia parviflora)*; patchy-canopy and moderate plant diversity.
- Moist Shivalik Sal Forest (hereafter Moist Forest): Dense forests on moist, north facing slopes of Shivaliks; dominated by *S. robusta*; more uniform-canopy and relatively less plant diversity.
- Dry Shivalik Sal Forest (hereafter Hill Forest): Open wooded slopes with scattered *S. robusta* and *Anogeissus latifolia*, and extensive grassy slopes and highest plant species diversity.

Precipitation is highest in the Moist forest followed by the Dry forest, and lowest in the Hill forest. The study area spans two north Indian states, Uttarakhand and Uttar Pradesh and lies in three forest administrative units, 1) Rajaji Tiger Reserve, 2) Shivalik Forest Division and 3) Dehradun Forest Division. This classification represents the protection level, management practices, and resource extraction patterns by communities. Study sites within Rajaji Tiger Reserve receive the highest level of protection and management interventions with restrictions on resource extraction (Fig. 1). Whereas sites within Dehradun and Shivalik Forest divisions experience relatively less protection along with nominal restriction on extraction of forest resources.

### 2.2 Sampling design

A total of 30 sampling cells of one square-kilometer were selected across the landscape with 10 cells in each forest type. Cell size was found optimum to capture the gradient of disturbances that could potentially influence vegetation characteristics yet conserve the forest composition and structure to enable comparisons. We ensured that the neighborhood of sampling cells was floristically alike to avoid edge effects confounding our inferences on habitat-disturbance relationships. Cells were first shortlisted using management plans and later selected based on reconnaissance surveys. Each cell had nine sampling points separated from the nearest point by >250 m. A total of 269 (3 forest types × 10 grids × 9 sampling point; one grid had only 8 points) points were marked within the study area and were sampled during summer and winter months between 2009–2011.

### 2.3 Vegetation quantification

At each sampling point, we quantified vegetation structure and composition of tree and shrub layers within 10 m and 5 m concentric circular plots, respectively. Woody plants ≥20 cm girth at breast height (GBH) were considered trees, and plants < 20 cm GBH were considered as shrubs. In a 10 m radius circular plot, we recorded the species identity, GBH, crown cover and height (bole and crown height) of each tree. Canopy cover was recorded only at the center of the plot using a convex spherical densitometer. For shrubs, both native and non-native, we recorded the species identity, number of individuals, height and canopy spread in two perpendicular diameters within the 5 m radius circular plot.

### 2.4 Anthropogenic disturbance

We quantified disturbance at each sampling point within a 20 m radius plot, concentric to the vegetation plot. Here, we recorded the percentage of lopped trees, number of dung pats (a proxy for the intensity of livestock grazing), the number and width of human trails intersecting the plot, the number of cut trees (<20 cm GBH) and cut stumps (>20 cm GBH) and percent of area grazed. Majority of our study sites were within tiger reserve and therefore we safely assumed that the large cut trees represented past legal logging. Percentage of area grazed was ocularly estimated by observing signs of browsing on shrub and herb layer within the sampling point. To reduce observer bias in estimating lopping intensity and percentage of grazing, these measurements were done by a single author (MK).

### 2.5 Data analysis

We examined if disturbance variables differed between forest types using a Kruskal-Wallis multiple comparison test (Siegel and Castellan, 1988). To identify factors that can potentially influence disturbance gradients, we: a) used Spearman’s correlation to analyze the association between disturbance components and human proximity, controlling for forest types; and b) tested if disturbance components differed between protection levels using Welch’s t-test (Sokal and Rohlf, 1995). Human proximity for each cell was quantified from village and road layers in program ARCMAP v 10 (ESRI, 2011).

To reduce dimensionality yet identify dominant patterns of vegetation structure, we extracted synthetic variables (components) from tree and shrub data using Principal Component Analysis (PCA). Principal components with eigenvalues greater than one were used to represent vegetation structure while analyzing vegetation-disturbance relationships. Since our sampling units were points nested within cells, we expected them to be spatially autocorrelated. We handled this hierarchically structured sampling design analytically in a mixed-effect modeling framework (Schielzeth and Nakagawa, 2013) and investigated the effects of disturbance variables on a) vegetation structure components, b) tree diversity, and c) shrub diversity. We used disturbance variables and forest types as fixed effects and variation between cells as random intercepts. Of all the disturbance variables, we selected the one that showed considerable variation for further modeling. For each of these responses, we built candidate models reflecting hypothesized effects, and compared them using Information Theoretic approach (Burnham and Anderson, 2002). Adequate model specification was confirmed by visual inspection of the residuals (Bates et al., 2014).

To examine the effects of disturbances on forest age structure (regeneration), we computed regression slopes of log-transformed tree counts against mid-points of GBH classes. To account for inherent difference in GBH among tree species, which impeded direct comparison of GBH across species, we scaled the GBH of each tree by the maximum GBH of that species so that all GBH values range between 0 and 1. Near-zero slopes reflected stable/declining age structure with an equal proportion of old and young trees, while steeper negative slopes reflected growing age structure with relatively more young than old trees (higher regeneration). We modeled slope values on disturbance variables and forest types using linear modeling, segregating the analysis between preferred and non-preferred fodder species (following preference values of Edgaonkar 1995). We also carried out the analysis for two important and dominant tree species namely *S. robusta* (a preferred fodder species) and *Mallotus phillipensis* (non-preferred fodder species).

We examined the effect of disturbance variables and forest types on *L. camara* invasion, using a hurdle/conditional modelling approach (Zuur et al., 2009). This is a two-step process, whereby *L. camara* presence vs. absence was first modeled using binomial generalized linear models, and then, varying abundance of *L. camara* in ‘presence only’ points was modeled using general linear models (Fletcher, MacKenzie, & Villouta, 2005; Welsh, Cunningham, Donnelly, & Lindenmayer, 1996). All analyses were performed using program R v 3.0.2 (R Development Core Team, 2013) using packages lme4 (Bates et al., 2014), MuMIn (Barton, 2014). Graphical output were prepared using packages ggplot2 (Wickham, 2009), sjplots (Lüdecke, 2016), and effects (Fox, 2003).

## 3. Results

We recorded 2758 adult trees belonging to 54 species across three forest types. Overall tree composition was dominated by the overstory species *S. robusta* (35% relative abundance) and its associated understory tree *M. phillipensis* (28%), followed by less common understory species such as *Ehretia laevis* (7%) and *Holarrhena antidysenterica* (5%), and the overstory species *A. latifolia* (4%).

### 3.1 Anthropogenic disturbances

Hill forest had significantly higher lopping and grazing than the other forest types (See Table A1 Supporting Information). Moist forest had higher firewood and timber extraction compared to other forest types. Trail width and number of dung pats did not differ between forest types (Fig. 2). Lopping was negatively associated with distance from village (Lopping: ρ = −0.42, p<0.05). Firewood collection was positively related to village density (ρ =0.55, p<0.001). Firewood collection and selective logging were significantly different across protected and less protected sites (Firewood collection: t = 2.69, df = 10, p-value <0.05; Selective logging t = 3.54, df = 11, p-value <0.005). Lopping and grazing did not differ across protection level. None of the disturbance variables were associated with road density and distance from road.

**Figure 2:**
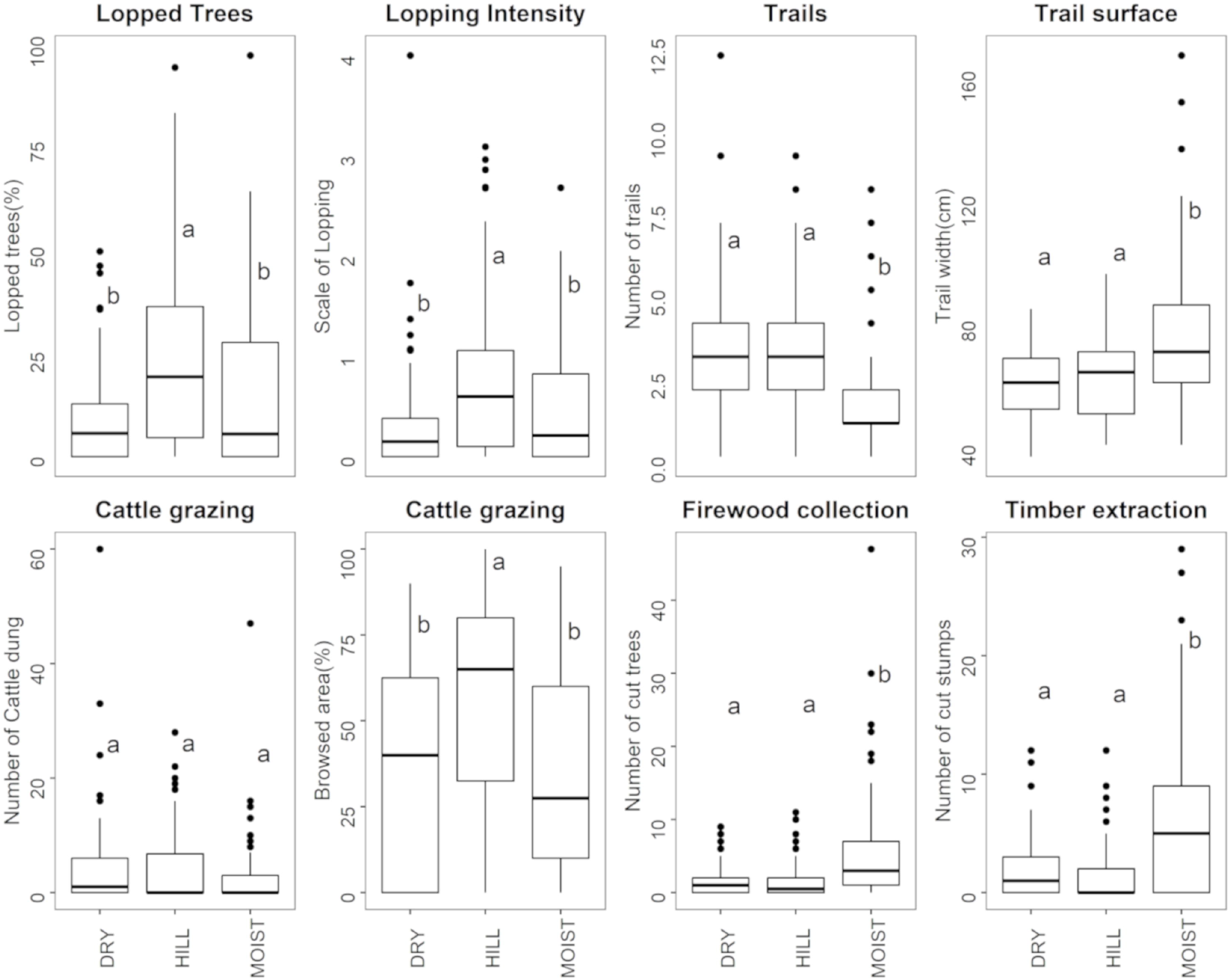
Statistical distribution of disturbance variables across forest types in the Shivalik region of the Himalayan Foothills, India. Variables with same letter code are not different from each other based on Tukey’s multiple comparison at p<0.05.

### 3.2 Vegetation characteristics

Moist forest differed structurally from other forest types, and was characterized by relatively tall, dense trees with large girths. However, floristic diversity was similar among forest types. Dry forest had highest density of exotic invasive shrub, *L. camara* (Table 1). Three principal components collectively explained 76% of variance in vegetation structure data (Table A2 Supporting Information). The first component represented mature forest features (higher values indicating taller and thicker trees with dense foliage) and explained 32% of the variance. The second component represented shrub biomass (higher values indicating tall and dense shrubs) and explained 24% of the variance. The third component represented secondary forest features (higher values indicating stands with dense canopy owing to high tree density rather than dense foliage) and explained 19% of the variance.

**Table 1:** Mean ± 1 SE (in parentheses) of vegetation structure and composition variables across three forest types in the Shivalik region of the Himalayan Foothills (India), estimated from 10 plots in each forest type.

|  | Dry Forest | Hill Forest | Moist Forest |
| --- | --- | --- | --- |
| **Vegetation Structure** |  |  |  |
| Tree density (per ha) | 278.14 (14.69) | 287.07 (16.08) | 406.92 (20.24) |
| Stand basal area (m<sup>2</sup> per ha) | 171.51 (10.55) | 143 (10.64) | 317.21 (10.87) |
| Tree height (m) | 15.85 (3.47) | 11.19 (1.72) | 20.67 (2.08) |
| Tree crown cover (m<sup>2</sup>) | 30.80 (6.61) | 21.78 (5.58) | 44.04 (17.32) |
| Canopy cover (%) | 65.39 (2.65) | 41.40 (2.72) | 87.00 (1.49) |
| Native shrub density (per m<sup>2</sup>) | 1.63 (0.11) | 0.21 (0.01) | 2.45 (0.24) |
| Native shrub cover (per m<sup>2</sup>) | 2.97 (2.11) | 1.18 (1.22) | 1.89 (1.48) |
| Native shrub height (m) | 1.20 (0.17) | 1.01 (0.13) | 1.19 (0.15) |
| Invasive *L. camara* cover (per m<sup>2</sup>) | 5.05 (1.03) | 1.99 (0.35) | 3.63 (0.54) |
| **Vegetation Composition** |  |  |  |
| Tree richness | 3.63 (1.41) | 4.18 (1.81) | 2.51 (1.22) |
| Tree diversity | 1.02 (0.04) | 1.66 (0.05) | 0.61 (0.05) |
| Shrub richness | 8.33 (0.73) | 8.13(0.98) | 8.13(0.70) |
| Shrub diversity | 1.94 (0.03) | 1.44 (0.05) | 1.29 (0.04) |

### 3.3 Vegetation structure – disturbance relationships

Among nine *a priori* hypotheses explaining mature forest features (vegetation PC 1), the model with timber extraction and forest type had maximum support (*wi*=0.72, see appendix A3) and explained 25% of the variation. Mature forest features decreased with increasing timber extraction (β= −0.20 ± 0.07) and were relatively higher in Moist Forest compared with other forest types (Fig. 3a). Out of 14 *a priori* hypotheses explaining shrub biomass (vegetation PC2), the model with grazing pressure had maximum support (*wi*=0.90, Table A3 in Supporting Information). Grazing pressure negatively influenced shrub biomass across forest types (β= −0.36 ± 0.08) and explained 13% of the variation (Fig. 3b). Comparison of 12 *a priori* hypotheses for explaining secondary forest features (vegetation PC3) showed maximum support for timber extraction, lopping and forest types (*wi*=0.71, Table A3 in Supporting Information), explaining 61% of the variation. Secondary forest features were positively associated with timber extraction (β= 0.11 ± 0.05), negatively associated with lopping (β= −0.22 ± 0.05) and were inherently different between forest types (Fig. 3c & 3d). These fixed effects explained a substantial proportion (44%) of the variation in secondary forest features.

**Figure 3:**
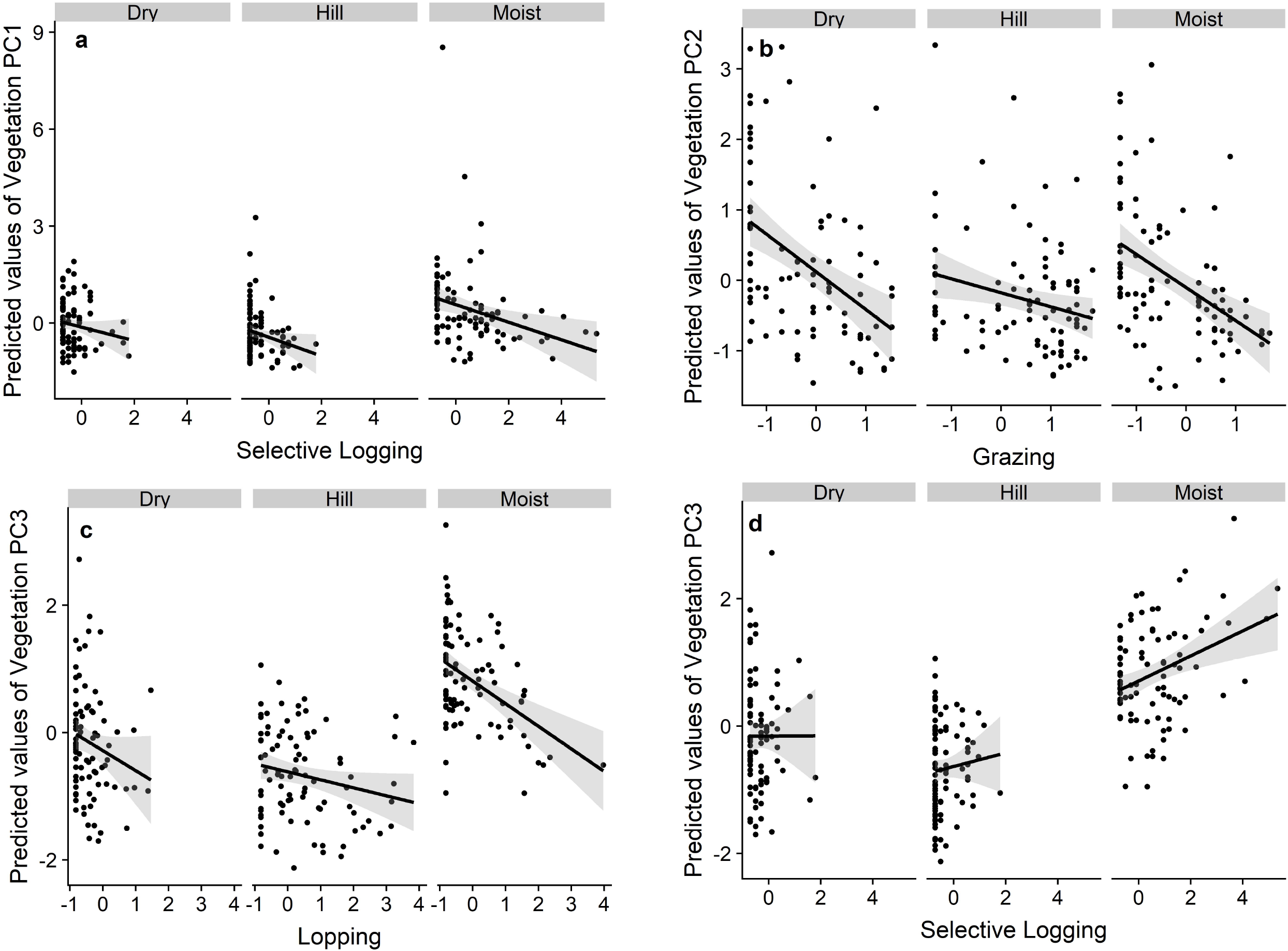
Responses of vegetation structural variables: (a) principle component 1 or mature forest features, (b) component 2 or shrub biomass, and (c & d) component 3 or secondary forest features, to anthropogenic disturbances such as selective logging, grazing and lopping across forest types in the Shivalik region of the Himalayan Foothills, India.

### 3.4 Floristic diversity – disturbance relationships

For tree diversity, the interactive effect of grazing pressure and forest types along with the additive effect of timber extraction obtained maximum support among the alternate hypotheses (*wi*=0.84, appendix A3) and explained 27% of the total variation (Fig.4a, 4b). Shrub richness was modeled against disturbance variables under the set of 14 *a priori* hypotheses, of which the model with an additive effect of grazing and forest type found maximum support (*wi*=0.53, appendix A3). Grazing reduced shrub richness across forest types (Fig. 4c) and this model explained 18% of the total variation. Among 15 *a priori* hypotheses for shrub diversity, the model with lopping pressure and forest types found maximum support (*wi*=0.70, appendix A3) and explained 60% of the total variation (Fig.4d).

**Figure 4:**
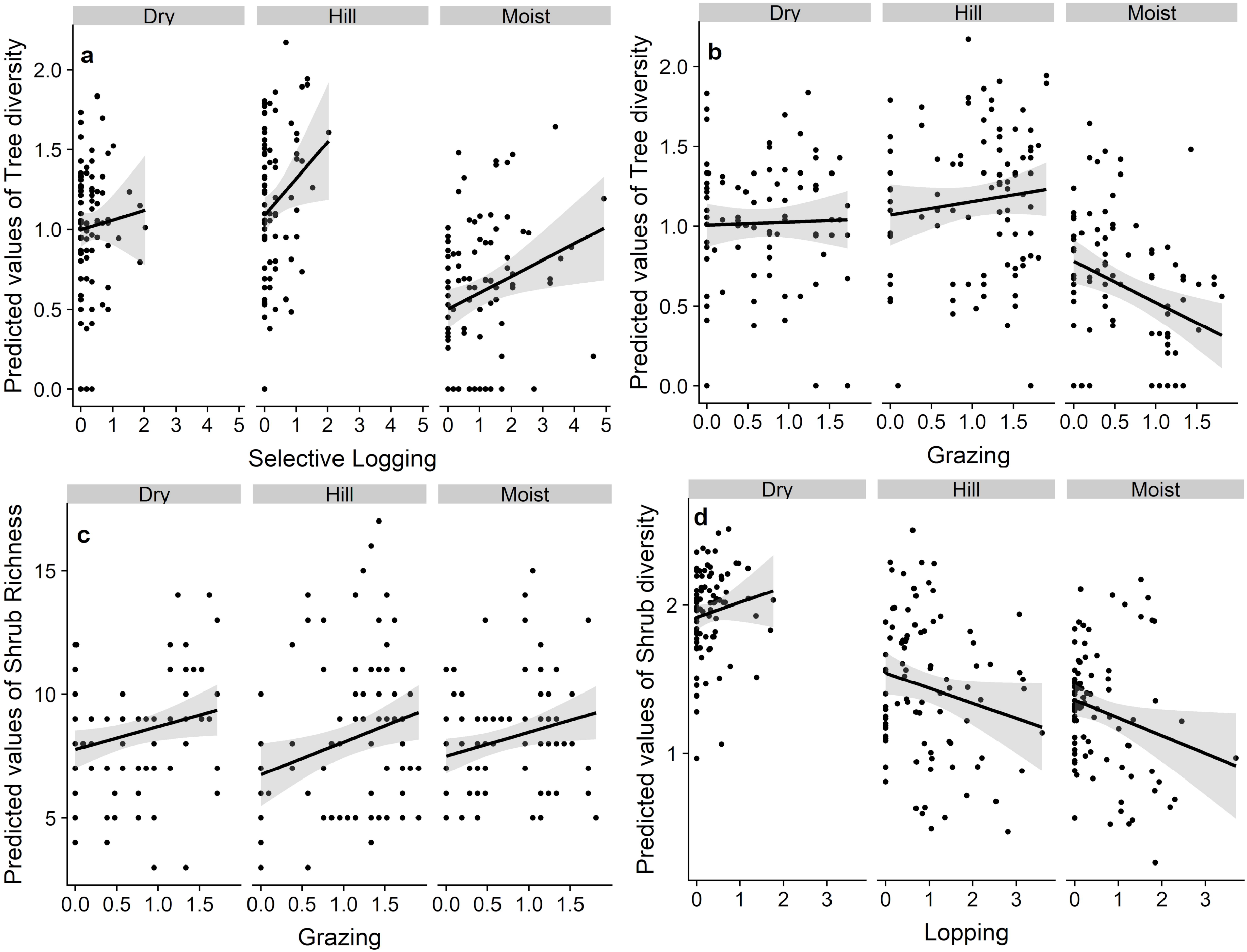
Responses of tree diversity (a & b) and shrub richness and diversity (c & d) to anthropogenic disturbances such as selective logging extraction, grazing and lopping across three forest types in the Shivalik region of the Himalayan Foothills, India.

### 3.5 Forest girth (age) distribution

Girth class distribution of trees in Dry and Hill forests (Fig. 5a) exhibited a typical inverse J shaped curve (Parthasarathy, 1999). However, there was an apparent deficit of individual trees in the smallest GBH class in Moist forest. Analysis of GBH slopes (log-counts regressed on GBH class) showed that the forest age structure was shaped by complex interactions between disturbance and forest types (Table A4 Supporting Information). Both selective logging and firewood collection had a negative association with the slopes of GBH, indicating steeper slopes of regeneration and thereby higher regeneration. The effect of lopping differed across preferred and non-preferred fodder tree species (Fig. 5b-5d). Lopping resulted in steeper slopes for preferred fodder tree species but shallower slopes for non-preferred fodder tree species and *M. phillipensis* (Fig. 5e). The interaction between selective logging and lopping also resulted in shallower slopes of GBH (positive values of model coefficient) for preferred fodder tree species (Fig. 5b). Forest types also differed in terms of girth class distribution. The Hill forest had the highest negative association with the GBH slopes for the preferred and non-preferred fodder tree species indicating higher regeneration. Slopes of GBH for *S. robusta* had the maximum negative association with Moist Forest followed by Dry and Hill forest.

**Figure 5:**
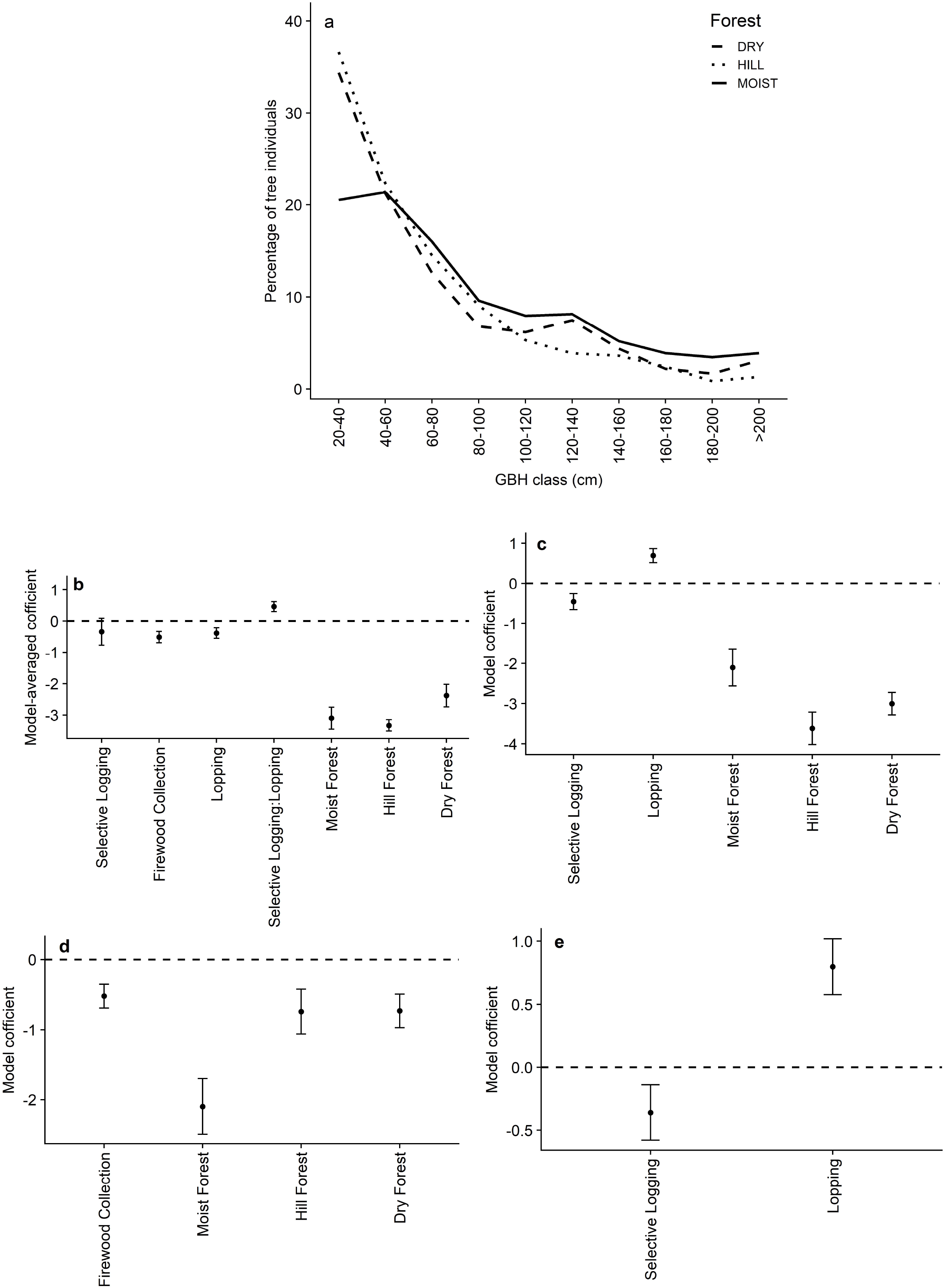
Frequency distribution of trees ((a); all woody plant with GBH >20 cm) compared across three forest types in the Shivalik region of the Himalayan Foothills, India. Influence of disturbance and forest types on GBH slope of (b) preferred fodder tree species (c) non-preferred tree species (d) *Shorea robusta* and (e) *Mallotus phillipensis*. Black dots represent the estimated values of the model parameter and error bars indicate standard error. Negative value of model coefficient implies steeper slopes (regenerating stand with a higher number of smaller GBH classes) whereas the positive value of model coefficients implies shallower slopes (arrested regeneration with lower number of small GBH classes).

### 3.6 Lantana camara invasion

Timber extraction and forest type best explained the presence of *L. camara* (*wi*=0.97, appendix A3). Timber extraction explained 25% of the variation in *L. camara*’s presence whereas the random effect of cells explained 52% variation. *L. camara* presence increased in Dry and Hill forest plots that were subjected to timber extraction in the past, whereas, the trend was opposite in Moist forest (Fig. 6a). Best-fit models showed high predictive accuracy for *L. camara* presence (Area under ROC curve=0.88). *L. camara*’s abundance was explained by the quadratic term of the percentage of grazing and forest type (*wi*=0.88, appendix A3). Models explaining *L. camara* abundance in invaded plots showed that their cover peaked at moderate levels of grazing pressure across forest types (Fig. 6b).

**Figure 6:**
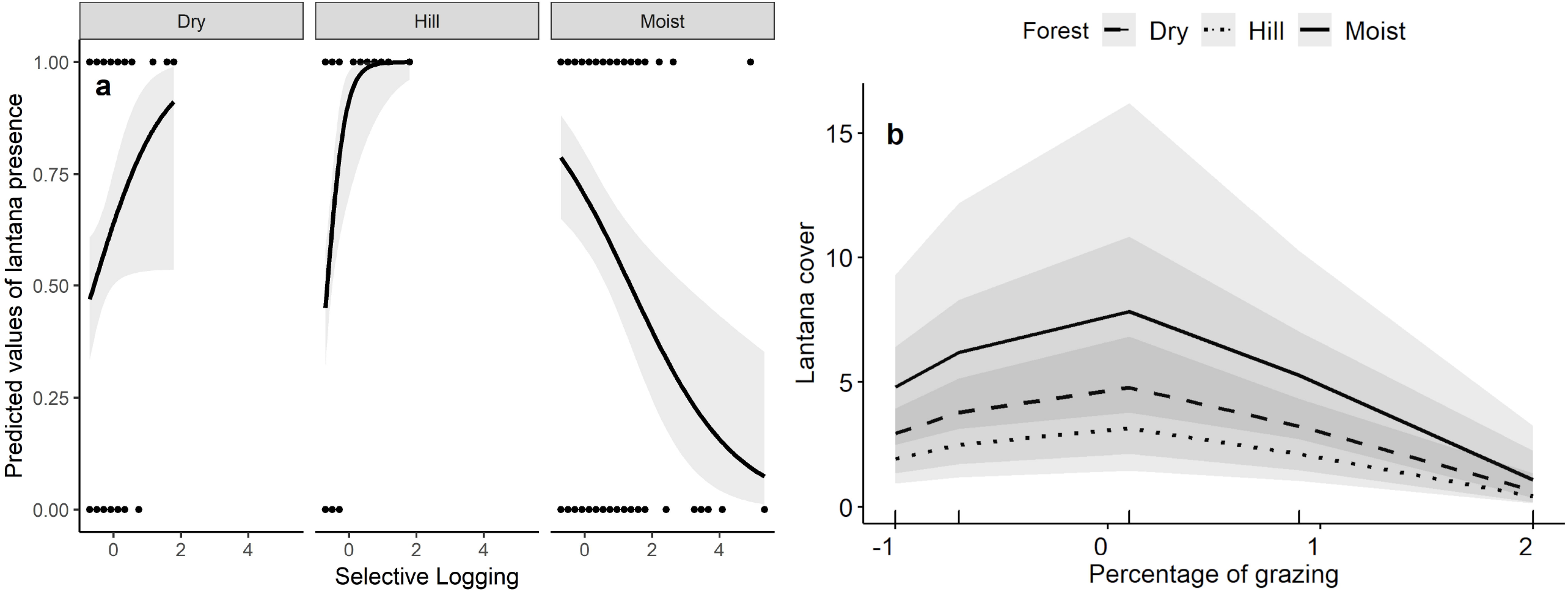
Responses of *Lantana camara*, an exotic invasive shrub, (a) presence and (b) abundance in invaded plots to disturbances such as selective logging and grazing pressure in the Shivalik region of the Himalayan Foothills, India.

## 4. Discussion

Comprehensive understanding of the impacts of CADs on forests is critical to sustaining tropical biodiversity and ecosystem services. Our study demonstrates that even small-scale continuous harvesting of forest resources can affect the structure, diversity, and regeneration of forest ecosystems, especially those with a history of large-scale disturbance (Chazdon, 2003). We also found legacy effects of 30-year-old selective logging as forestry operations on vegetation structure, diversity and invasion by *L. camara* in the sub-tropical Shivalik forests of Himalayan foothills. Even though selective logging has ceased in these tracts, ongoing CADs continue to influence forest structure and composition by shifting recovery trajectory. For example, gaps created by past selective logging facilitated invasion of the exotic shrub *L. camara* in drier forests. Furthermore, moderate levels of grazing pressure facilitated expansion of *L. camara* within colonized patches, which highlights the synergistic effects of past selective logging and present CADs on forest dynamics, especially in drier forests. As elaborated in the following sections, an important insight from this study was the differential response of certain vegetation parameters across forest types. Thus, we urge conservation practitioners to take caution in making generalizations about the impacts of disturbances on tropical forest dynamics.

### 4.1 Disturbance proxies

Applied ecological studies frequently use proxies for anthropogenic disturbances to examine the latter’s effects on ecological parameters and processes of interest, such as regeneration (Popradit et al., 2015), on the premise that these proxies are correlated with cryptic and difficult to measure CADs. Studies have used proximity to roads and villages, density of human and livestock and size of villages as indicators of disturbances (Karanth et al., 2006; Ribeiro-Neto et al., 2016). Although these secondary or remotely sensed indicators can allow rapid quantification of potential habitat degradation, few have examined whether they are accurate proxies for CAD impacts. Our study demonstrated that village density was a good indicator of lopping and firewood collection. However, illicit timber extraction was not explained by village density but by protection status. Higher protection discouraged villagers to harvest resources in large quantities from Tiger Reserve compared to less protected Reserve Forests. Interestingly, none of the disturbance variables were correlated with road density in our study area. Villagers and forest dwellers used dirt trails rather than roads for harvesting forest resources. Moreover, timber poachers might not use major road networks to transport illegally extracted timber. Our results support the value of increased protection in preventing habitat degradation from large-scale disturbances, while also emphasizing the importance of field-validation of disturbance indicators.

### 4.2 Influence of disturbance on vegetation structure

In the current study, 30-year-old selective logging was associated with decreased mature forest characteristics (greater tree height, GBH, basal area and crown cover) and increased secondary forest characteristics (greater tree density and percent canopy cover) across all forest types. This result resonates with similar studies conducted in the rainforest of Malaysia (Okuda et al., 2003), evergreen moist forest of Western Ghats (Pélissier et al., 1998) and moist forest of Uganda (Plumptre, 1996) where the forest still carries the legacy of past logging even decades after its cessation. In particular, structural features such as crown cover (Okuda et.al. 2003), height and basal area (Hawthorne et al., 2012; Plumptre, 1996) were significantly lower in logged tropical forest across countries. Our study area has not experienced any mechanized logging for at least three decades since the enactment of the Forest Conservation Act (1980). Thus, our results along with others’ highlight that forests may take decades to completely recover from large-scale logging that is significant in the context of proposed revisions of India’s National Forest Policy bill bringing back industrial forestry (Ministry of Environment, Forest and Climate Change., 2018).

We found that selective logging facilitated *L. camara* colonization especially in drier forests. The already sparse canopy of Hill and Dry forests with low soil moisture regimes might have made them more prone to *L. camara* invasion following disturbance from selective logging. However, we found the opposite effect in Moist forest, which could be attributed to relatively high canopy cover (76.7-96.5%, Table 1) and soil-moisture content. For instance, we found that areas in the Vindhyan highlands with more than 63% canopy cover and moderate soil moisture (7.5-30%) did not face *L. camara* invasion (Sharma and Raghubanshi, 2010).

Pastoral disturbances were negatively associated with secondary forest characteristics and shrub biomass. The negative association between lopping and secondary forest characteristics might result from a preference for large trees with bigger crowns for fodder extraction. Alternatively, partial lopping could increase GBH, another mature forest characteristic, by increasing the photosynthetic rate in remaining leaves (Bhat et al., 1995). However, this result could also indicate causation because of the negative consequences of lopping on sapling regeneration and establishment. Lopping poses continuous pressure on preferred trees resulting in loss of leafy biomass (Bhat et al., 1995) and ultimately lowering fruit and seed set (Sinha and Bawa, 2002). Lopping also reduces the amount of litter falling on ground, thereby reducing nutrients leaching to the soil (Melkania and Ramnarayan, 1998) and exposes the forest floor to higher temperatures. All these factors can suppress natural regeneration and secondary forest features such as higher tree density and canopy cover. This study also found a negative association of lopping on the regeneration of non-preferred fodder tree species including *M. phillipensis* (see Fig. 5c & 5e). Although lopping promoted regeneration of preferred fodder trees, the interaction of lopping and selective logging inhibited regeneration (Fig. 5b).

Livestock grazing and associated soil disturbance has been shown to favor non-native over native shrubs in temperate Australian grasslands (McIntyre and Lavorel, 1994) and in the Douglas-fir forest of the western United States (Zimmerman and Neuenschwander, 1984). In our study, moderate levels of livestock grazing also led to increased *L. camara* abundance. Although we could not investigate the effect of *L. camara* on native tree and shrub species due sparse data, we expect that such invasions of woody shrub species could lead to gradual modification of structural, compositional and functional characteristics of vegetation communities in Shivalik as observed in dry tropical forests in India (Sharma et al., 2005; Sundaram and Hiremath, 2012). These patterns have been observed elsewhere; increased *L. camara* cover in Australian’s Sclerophyll forests led to decreased native shrub and tree densities (Gooden *et al*., 2009). Plant species richness in the tropical dry forests of Queensland decreased with increasing *L. camara* density, and *L. camara* invaded wet Sclerophyll forest of Australia which had fewer plant species than the non-invaded areas (Fensham *et al*., 1994; Gooden *et al*., 2009).

### 4.3 Floristic diversity

Our results indicated that past selective logging favored tree diversity across forest types, but grazing pressure suppressed tree diversity in moist forests. Past selective logging could enhance overall tree species diversity by removing common canopy species and facilitating the colonization of pioneer species. This relationship largely holds true for low-intensity logging (Parrotta et al., 2002), areas with high local species richness, and viable seed disperser populations (Chazdon, 2003). However, high-intensity CADs can lead to poor germination due to severe soil compaction, competition by non-native invasive species (Collier et al., 2002) or high mortality of seedlings owing to browsing by livestock. Wider trails in Moist forest indicates that these areas may be used at higher frequency than the actively protected drier forest in the study area. Soil compaction could result in poor infiltration and greater runoff, thereby reducing seed germination (Chazdon, 2003). Additionally, steeper curves of tree and shrub diversity due to CADs in Moist forest could be attributed to higher rates of seedling damage from pathogens and pests (Brenes-Arguedas et al., 2009). Shrub cover facilitates tree seedling establishment by promoting seed rain and providing optimum nutrient and microclimatic conditions (Holl, 2002; Vieira et al., 1994). Severe declines in shrub height and cover in Moist forest (Fig. 3b) could be associated with declines in tree and shrub diversity.

## 5. Conclusion

We demonstrate the direct and indirect impacts of CAD on vegetation structure, diversity, regeneration and invasion. With a growing human population and increased consumption, we expect these pressures on forests in Himalayan Foothills to grow in the future. Therefore, we suggest that following is warranted: (i) it is time to acknowledge the negative consequences of increasing small-scale harvest on forest ecosystems, (ii) the protection status of the reserved forest should be upgraded to reduce illicit timber extraction, (iii) additional research on alternative livelihood options to reduce the dependency of rural communities on forests is much needed, (iv) forest managers should consider actions to enhance recruitment of tree species facing severe regeneration failure, and (v) the control of non-native invasive species should be actively enforced per existing afforestation policies (i.e., Compensatory afforestation fund management and planning authority (CAMPA) in India). Ignoring chronic and seeming small-scale cryptic disturbances can result in further loss of biodiversity and ecosystem services in forests with high conservation value through complex and interacting mechanisms.

## Author’s Contribution

D.M. and P.S. were the investigators of the project that funded this study. M.K., S.D. and G.S.R. conceived this paper. M.K collected field data and analyzed it with inputs from SD. MK wrote the paper with contributions from all authors in methodology, interpretation and editing. All authors critically assessed and approved the final version of the manuscript.

## Acknowledgements

This study was funded by the Wildlife Institute of India’s grant-in-aid. We acknowledge support received from Raman Kumar, Soumya Prasad, Rajah Jayapal, Qamar Qureshi, Trevor Price for designing study and analyzing data. We thank Uttarakhand and Uttar Pradesh forest departments for research permits and logistical support. We are grateful to Inam Ali, Kuldeep and Bhura for assisting in field. We thank Ninad Mungi for helping with study area map. We are grateful to Liba Pejchar for providing useful comments on the manuscript.

## Supporting information

**Additional details on disturbance variation across forest types, loading of principal components and model specification of candidate models. See Table A1, A2, A3 and A4**.

## Notes

### Competing Interest Statement

The authors have declared no competing interest.

