## Supplemental Table 1 for "Disentangling the effects of past logging and ongoing cryptic anthropogenic disturbance on vegetation structure and composition in Himalayan Foothills, India"

| Disturbance variables | | | Dry Forest | |  | | Hill Forest |  | Moist Forest | |
| --- | --- | --- | --- | --- | --- | --- | --- | --- | --- | --- |
|  | Average | | | Range | | Average | Range | Average | | Range |
| Lopping Intensity | | 0.3 | | (0‐4) | 0.8 | | (0‐3) | 0.5 | | (0‐2.7) |
| Lopped Trees (%) | | 9.7 | | (0‐55) | 24.8 | | (0‐93) | 14.8 | | (0‐95) |
| Number of Trails | | 2.7 | | (0‐12) | 3.0 | | (0‐9) | 2.0 | | (0‐8) |
| Trail width (cm) | | 54.9 | | (0‐84) | 59.4 | | (0‐95) | 62.3 | | (0‐165) |
| Number of cattle dung | | 4.2 | | (0‐60) | 4.4 | | (0‐28) | 2.7 | | (0‐47) |
| Number of cut trees (20- 40cm GBH class) | | 1.3 | | (0‐9) | 1.5 | | (0‐11) | 5.6 | | (0‐47) |
| Number of cut trees (>40cm GBH class) | | 2.1 | | (0‐12) | 1.8 | | (0‐12) | 6.3 | | (0‐29) |
| Area grazed (%) | | 37.4 | | (0‐90) | 55.2 | | (0‐100) | 34.6 | | (0‐95) |

**Table A1**: Mean & range of disturbance variables across forest types. Range refers to the minimum and maximum value attained by a disturbance variable in a given forest type across all plots.
