## Supplemental Table 2 for "Disentangling the effects of past logging and ongoing cryptic anthropogenic disturbance on vegetation structure and composition in Himalayan Foothills, India"

**Table A2**: Metric loadings for the first three principal components

|  | PC1 | PC2 | PC3 |
| --- | --- | --- | --- |
| Mature Forest Features |  |  |  |
| Tree GBH | 0.92 | 0.07 | -0.16 |
| Tree Crown Cover | 0.74 | 0.24 | -0.14 |
| Tree Height | 0.74 | 0.01 | 0.33 |
| Shrub Volume |  |  |  |
| Shrub Cover | 0.20 | 0.83 | 0.10 |
| Shrub Height | 0.02 | 0.92 | 0.01 |
| Secondary Forest Features |  |  |  |
| Percentage Canopy Cover | 0.41 | 0.28 | 0.73 |
| Tree Density | -0.35 | -0.06 | 0.81 |
