## Supplemental Table 3 for "Disentangling the effects of past logging and ongoing cryptic anthropogenic disturbance on vegetation structure and composition in Himalayan Foothills, India"

**Table A3.** **A priori hypothesized models built to explain the effects of forest type, chronic and acute anthropogenic disturbance variable on vegetation structure (Vegetation PC1, Vegetation PC2 and Vegetation PC3), vegetation diversity (Tree richness, Shrub richness, tree diversity and shrub diversity and exotic invasive *Lantana camara*’s presence and abundance in the Shivalik region of Himalayan foothills, India. Models are ranked by scores of Akaike's information criterion adjusted for small sample size (AIC). Columns include the covariates used in the model including intercept, AIC score, distance from the lowest AIC (ΔAIC) and Akaike's model weight (*wi*).**

| Response variable | Model | AIC | ∆AIC | | *w_i_* |
| --- | --- | --- | --- | --- | --- |
| Vegetation PC1 | Selective Logging+ Forest+ (1\|Plot) | 719.8 | 0 | | 0.72 |
|  | Forest+(1\|Plot) | 722.8 | 2.93 | | 0.17 |
|  | Selective Logging+ Lopping+ Forest+ (1\|Plot) | 725.2 | 5.32 | | 0.05 |
|  | Null Model | 726 | 6.21 | | 0.03 |
|  | Selective Logging x Forest+ (1\|Plot) | 726.5 | 6.71 | | 0.03 |
|  | Lopping + Forest + (1\|Plot) | 728 | 8.16 | | 0.01 |
|  | Lopping x Forest + (1\|Plot) | 733.2 | 13.38 | | 0.00 |
|  | Full Model | 735 | 15.18 | | 0.00 |
|  | Lopping x Forest + Selective Logging x Forest+(1\|Plot) | 737.4 | 17.59 | | 0.00 |
| Vegetation PC2 | Grazing+(1\|Plot) | 675.7 | 0 | | 0.90 |
|  | Grazing + Forest+(1\|Plot) | 680.2 | 4.54 | | 0.09 |
|  | Grazing + (Grazing^2^) +Forest+(1\|Plot) | 685.4 | 9.75 | | 0.01 |
|  | Grazing X Forest+(1\|Plot) | 687.2 | 11.53 | | 0.00 |
|  | Full Model | 691.2 | 15.49 | | 0.00 |
|  | Null Model | 692.2 | 16.51 | | 0.00 |
|  | Selective Logging+(1\|Plot) | 693 | 17.35 | | 0.00 |
|  | Forest+(1\|Plot) | 694.8 | 19.12 | | 0.00 |
|  | Selective Logging + Forest+(1\|Plot) | 695.2 | 19.52 | | 0.00 |
|  | Lopping+(1\|Plot) | 695.9 | 20.2 | | 0.00 |
|  | Lopping + Forest+(1\|Plot) | 699.1 | 23.48 | | 0.00 |
|  | Lopping + Selective Logging + Forest+(1\|Plot) | 699.6 | 23.9 | | 0.00 |
|  | Selective Logging X Forest+(1\|Plot) | 700.2 | 24.58 | | 0.00 |
|  | Lopping X Forest+(1\|Plot) | 703.6 | 27.94 | | 0.00 |
| Vegetation PC3 | Selective Logging + Lopping + Forest+(1\|Plot) | 570.2 | 0 | 0.71 | |
|  | Lopping + Firewood Collection + Selective Logging + Forest+(1\|Plot) | 573.9 | 3.64 | | 0.12 |
|  | Lopping X Forest +Selective Logging+(1\|Plot) | 574.3 | 4.09 | | 0.09 |
|  | Selective Logging X Forest+ Lopping+(1\|Plot) | 576 | 5.71 | | 0.04 |
|  | Forest + (1\|Plot) | 577 | 6.78 | | 0.02 |
|  | Selective Logging + Forest+(1\|Plot) | 578.6 | 8.33 | | 0.01 |
|  | Selective Logging + Firewood Collection+ Forest +(1\|Plot) | 582.5 | 12.27 | | 0.00 |
|  | Selective Logging X Forest+(1\|Plot) | 583.8 | 13.54 | | 0.00 |
|  | Selective Logging X Forest + Firewood Collection X Forest+(1\|Plot) | 594 | 23.76 | | 0.00 |
|  | Selective Logging+(1\|Plot) | 596.9 | 26.68 | | 0.00 |
|  | Null Model | 597.4 | 27.14 | | 0.00 |
|  | Selective Logging + Firewood Collection+(1\|Plot) | 598.4 | 28.19 | | 0.00 |
| Tree richness | Selective Logging +Grazing X Forest | 956.5 | 0 | | 0.60 |
|  | Selective Logging + Forest | 958.9 | 2.44 | | 0.18 |
|  | Selective Logging X Forest | 959.9 | 3.45 | | 0.11 |
|  | Selective Logging + Grazing + Forest | 960.8 | 4.3 | | 0.07 |
|  | Full Model | 962.9 | 6.41 | | 0.02 |
|  | Grazing X Forest | 963.7 | 7.24 | | 0.02 |
|  | Forest | 966.5 | 10.04 | | 0.00 |
|  | Lopping X Forest | 966.8 | 10.3 | | 0.00 |
|  | Null Model | 1001.3 | 44.8 | | 0.00 |
|  | Selective Logging | 1003.3 | 46.79 | | 0.00 |
| Shrub Richness | Grazing+(1\|Plot) | 1225.6 | 0 | | 0.53 |
|  | Grazing + Forest+(1\|Plot) | 1228.6 | 3.01 | | 0.12 |
|  | Null Model | 1228.9 | 3.25 | | 0.11 |
|  | Selective Logging + Grazing + Forest +(1\|Plot) | 1230.6 | 4.97 | | 0.04 |
|  | Grazing + (Grazing^2^) +Forest+(1\|Plot) | 1230.6 | 5 | | 0.04 |
|  | Lopping+(1\|Plot) | 1230.9 | 5.25 | | 0.04 |
|  | Selective Logging+(1\|Plot) | 1230.9 | 5.25 | | 0.04 |
|  | Selective Logging X Forest+(1\|Plot) | 1231.6 | 5.99 | | 0.03 |
|  | Grazing X Forest+(1\|Plot) | 1232.2 | 6.6 | | 0.02 |
|  | Forest+(1\|Plot) | 1232.6 | 7.03 | | 0.02 |
|  | Lopping + Forest+(1\|Plot) | 1234.6 | 9.01 | | 0.01 |
|  | Selective Logging + Forest+(1\|Plot) | 1234.6 | 9.03 | | 0.01 |
|  | Full Model | 1235.3 | 9.69 | | 0.00 |
|  | Lopping X Forest+(1\|Plot) | 1237.3 | 11.68 | | 0.00 |
| Tree diversity | Selective Logging + Grazing X Forest | 98.5 | 0 | | 0.84 |
|  | Selective Logging + Forest | 304.2 | 5.67 | | 0.05 |
|  | Selective Logging + Grazing + Forest | 304.7 | 6.16 | | 0.04 |
|  | Full Model | 304.8 | 6.25 | | 0.04 |
|  | Selective Logging X Forest | 306.5 | 7.93 | | 0.02 |
|  | Grazing X Forest | 307.4 | 8.91 | | 0.01 |
|  | Lopping X Forest | 309.6 | 11.1 | | 0.00 |
|  | Forest | 312.4 | 13.86 | | 0.00 |
|  | Null Model | 377.1 | 78.6 | | 0.00 |
|  | Selective Logging | 378.5 | 79.93 | | 0.00 |
| Shrub Diversity | Lopping + Forest + (1\|Plot) | 219.7 | 0 | | 0.70 |
|  | Forest + (1\|Plot) | 221.8 | 2.07 | | 0.25 |
|  | Grazing + Forest + (1\|Plot) | 226.9 | 7.25 | | 0.02 |
|  | Lopping X Forest + (1\|Plot) | 227.6 | 7.95 | | 0.01 |
|  | Selective Logging + Forest + (1\|Plot) | 228.5 | 8.83 | | 0.01 |
|  | Selective Logging X Forest + (1\|Plot) | 229.5 | 9.81 | | 0.01 |
|  | Lopping + (1\|Plot) | 231.2 | 11.47 | | 0.00 |
|  | Grazing+ (Grazing^2^) + Forest+(1\|Plot) | 231.3 | 11.65 | | 0.00 |
|  | Selective Logging + Grazing + Forest+(1\|Plot) | 233.6 | 13.94 | | 0.00 |
|  | Grazing X Forest + (1\|Plot) | 234.3 | 14.65 | | 0.00 |
|  | Null Model | 235.4 | 15.69 | | 0.00 |
|  | Grazing + (1\|Plot) | 240.4 | 20.71 | | 0.00 |
|  | Full Model | 240.5 | 20.79 | | 0.00 |
|  | Selective Logging + (1\|Plot) | 241.7 | 22.06 | | 0.00 |
|  | Grazing + (Grazing^2^) + (1\|Plot) | 244.4 | 24.75 | | 0.00 |
| Lantana's Presence | Selective Logging X Forest+(1\|Plot) | 301.2 | 0 | | 0.98 |
|  | Null Model | 310.6 | 9.4 | | 0.01 |
|  | Selective Logging+(1\|Plot) | 311.9 | 10.72 | | 0.01 |
|  | (Firewood Collection^2^) +Forest + Selective Logging+(1\|Plot) | 313.5 | 12.35 | | 0.00 |
|  | Lopping + Forest+(1\|Plot) | 314.1 | 12.87 | | 0.00 |
|  | Forest+(1\|Plot) | 314.6 | 13.36 | | 0.00 |
|  | Lopping + Selective Logging + Forest+(1\|Plot) | 315.3 | 14.15 | | 0.00 |
|  | Selective Logging + Forest+(1\|Plot) | 315.9 | 14.69 | | 0.00 |
|  | Lopping X Forest+(1\|Plot) | 316.1 | 14.95 | | 0.00 |
|  | (Lopping^2^) + Forest + Selective Logging+(1\|Plot) | 317.8 | 16.66 | | 0.00 |
|  | Full Model | 318.1 | 16.94 | | 0.00 |
|  | Firewood Collection X Forest + Selective Logging+(1\|Plot) | 319.2 | 18.05 | | 0.00 |
|  | Grazing X Forest + Selective Logging+(1\|Plot) | 321.9 | 20.69 | | 0.00 |
| Lantana's abundance | Firewood Collection X Forest+(Grazing^2^) + (1\|Plot) | 1072.2 | 0 | | 0.78 |
|  | Firewood Collection X Forest+(1\|Plot) | 1076 | 3.78 | | 0.12 |
|  | Firewood Collection X Forest + Lopping+(1\|Plot) | 1076.6 | 4.37 | | 0.09 |
|  | (Grazing2) + Forest + (1\|Plot) | 1083.1 | 10.86 | | 0.00 |
|  | Lopping X Forest + (1\|Plot) | 1084.1 | 11.81 | | 0.00 |
|  | Lopping X Forest + Firewood Collection+(1\|Plot) | 1084.3 | 12.07 | | 0.00 |
|  | Grazing X Forest+(1\|Plot) | 1085 | 12.78 | | 0.00 |
|  | (Firewood Collection^2^) + Forest+(Grazing^2^) + (1\|Plot) | 1086.3 | 14.06 | | 0.00 |
|  | Selective Logging X Forest+(1\|Plot) | 1086.5 | 14.25 | | 0.00 |
|  | Forest+(1\|Plot) | 1086.8 | 14.57 | | 0.00 |
|  | Lopping + Forest+(1\|Plot) | 1087 | 14.74 | | 0.00 |
|  | Lopping + Firewood Collection + Forest+(1\|Plot) | 1087.1 | 14.83 | | 0.00 |
|  | Lopping+(Lopping^2^) + Forest+(1\|Plot) | 1087.5 | 15.3 | | 0.00 |
|  | Grazing + Firewood Collection + Forest+(1\|Plot) | 1087.7 | 15.47 | | 0.00 |
|  | Selective Logging + Forest+(1\|Plot) | 1088 | 15.71 | | 0.00 |
|  | Lopping + Selective Logging + Forest+(1\|Plot) | 1088.1 | 15.89 | | 0.00 |
|  | Firewood Collection+ (Firewood Collection^2^) + Forest + (1\|Plot) | 1089.1 | 16.83 | | 0.00 |
|  | Null Model | 1093.6 | 21.32 | | 0.00 |
|  | Grazing+(1\|Plot) | 1093.7 | 21.5 | | 0.00 |
|  | Firewood Collection+(1\|Plot) | 1094.1 | 21.83 | | 0.00 |
|  | Lopping+(1\|Plot) | 1094.4 | 22.14 | | 0.00 |
|  | (Selective Logging^2^) + Forest+(1\|Plot) | 1096.6 | 24.39 | | 0.00 |
