## Supplemental Table 4 for "Disentangling the effects of past logging and ongoing cryptic anthropogenic disturbance on vegetation structure and composition in Himalayan Foothills, India"

**Table A4.** **A priori hypothesized models built to explain the effects of forest type, chronic and acute anthropogenic disturbance variable on the regeneration expressed as GBH of slope for preferred, non-preferred tree species, *Shorea robusta* and *Mallotus phillipensis* in the Shivalik region of the Himalayan Foothills, India. Models are ranked by scores of Akaike's information criterion adjusted for small sample size (AICc). Columns include the covariates used in the model including intercept, AICc score, distance from the lowest AICc (ΔAICc) and Akaike's model weight (*wi*).**

| Response variable | |  | | | AICc | ∆AICc | *w_i_* |
| --- | --- | --- | --- | --- | --- | --- | --- |
| Slope of GBH of Preferred Fodder tree species | | | Firewood collection+Forest | | 58.70 | 0.00 | 0.33 |
|  | | | Selective Logging X Lopping | | 59.00 | 0.28 | 0.29 |
|  | | | Selective Logging X Lopping + Forest | | 61.30 | 2.56 | 0.09 |
|  | | | Firewood Collection + Lopping + Forest | | 61.90 | 3.20 | 0.07 |
|  | | | Selective Logging | | 62.90 | 4.13 | 0.04 |
|  | | | Firewood Collection X Grazing | | 62.90 | 4.18 | 0.04 |
|  | | | Selective Logging + Lopping | | 63.10 | 4.32 | 0.04 |
|  | | | Selective Logging + Lopping + Forest | | 63.60 | 4.82 | 0.03 |
|  | | | Selective Logging + Grazing + Forest | | 63.80 | 5.06 | 0.03 |
|  | | | Full Model | | 66.00 | 7.24 | 0.01 |
|  | | | Selective Logging X Grazing | | 66.50 | 7.73 | 0.01 |
|  | | | Selective Logging X Forest | | 66.80 | 8.01 | 0.01 |
|  | | | Lopping + Forest | | 66.90 | 8.19 | 0.01 |
|  | | | Lopping X Forest + Selective Logging | | 68.10 | 9.37 | 0.00 |
|  | | | Null Model | | 69.30 | 10.51 | 0.00 |
| Slope of GBH of Non-Preferred Fodder tree species | Selective Logging+Lopping+Forest | | | | 64.4 | 0 | 0.42 |
|  | Lopping X Forest+Selective Logging | | | | 66 | 1.68 | 0.18 |
|  | Lopping+Forest | | | | 66.6 | 2.22 | 0.14 |
|  | Selective Logging+Lopping | | | | 67.3 | 2.93 | 0.10 |
|  | Selective Logging X Lopping+Forest | | | | 68.3 | 3.95 | 0.06 |
|  | Null Model | | | | 70 | 5.63 | 0.03 |
|  | Full Model | | | | 70.1 | 5.76 | 0.02 |
|  | Firewood Collection+Lopping+Forest | | | | 70.2 | 5.82 | 0.02 |
|  | Selective Logging X Lopping | | | | 70.4 | 6.08 | 0.02 |
|  | Selective Logging+Grazing+Forest | | | | 72.3 | 7.92 | 0.01 |
|  | Selective Logging | | | | 72.5 | 8.17 | 0.01 |
|  | Selective Logging X Grazing | | | | 74.7 | 10.36 | 0.00 |
|  | Firewood Collection X Grazing+Forest | | | | 76.7 | 12.38 | 0.00 |
|  | Firewood Collection+Forest | | | | 77.1 | 12.72 | 0.00 |
|  | Selective Logging X Forest | | | | 82.8 | 18.43 | 0.00 |
| Slope of GBH of *Shorea robusta* | Firewood Collection+Forest | | | | 57.40 | 0.00 | 0.67 |
|  | Firewood Collection + Lopping+Forest | | | | 60.80 | 3.34 | 0.13 |
|  | Firewood Collection X Grazing+Forest | | | | 60.80 | 3.41 | 0.12 |
|  | Selective Logging + Lopping+Forest | | | | 64.90 | 7.45 | 0.02 |
|  | Selective Logging + Grazing+Forest | | | | 65.00 | 7.56 | 0.02 |
|  | Selective Logging X forest | | | | 65.10 | 7.64 | 0.02 |
|  | Selective Logging | | | | 65.50 | 8.10 | 0.01 |
|  | Selective Logging X Lopping+Forest | | | | 66.80 | 9.40 | 0.01 |
|  | Lopping + Forest | | | | 67.00 | 9.53 | 0.01 |
|  | Full Model | | | | 67.80 | 10.34 | 0.00 |
|  | Selective Logging + Lopping | | | | 68.00 | 10.62 | 0.00 |
|  | Selective Logging X Grazing | | | | 69.80 | 12.38 | 0.00 |
|  | | | | Selective Logging X Lopping | 70.30 | 12.90 | 0.00 |
|  | | | | Lopping X Forest + Selective Logging | 72.40 | 15.02 | 0.00 |
|  | | | | Null Model | 80.00 | 22.59 | 0.00 |
| Slope of GBH of *Mallotus phillipensis* | | | | Selective Logging+Lopping | 77.70 | 0.00 | 0.65 |
|  | | | | Lopping X Forest + Selective Logging | 79.80 | 2.07 | 0.23 |
|  | | | | Lopping + Forest | 83.40 | 5.73 | 0.04 |
|  | | | | Selective Logging + Lopping | 84.00 | 6.32 | 0.03 |
|  | | | | Lopping + Forest | 84.90 | 7.16 | 0.02 |
|  | | | | Null Model | 85.30 | 7.60 | 0.01 |
|  | | | | Selective Logging | 86.40 | 8.71 | 0.01 |
|  | | | | Selective Logging X Grazing | 86.40 | 8.74 | 0.01 |
|  | | | | Firewood Collection X Grazing+Forest | 87.00 | 9.27 | 0.01 |
|  | | | | Full Model | 90.60 | 12.95 | 0.00 |
|  | | | | Selective Logging X Lopping | 91.60 | 13.89 | 0.00 |
|  | | | | Selective Logging + Grazing + Forest | 91.90 | 14.16 | 0.00 |
|  | | | | Firewood Collection + Forest | 93.00 | 15.26 | 0.00 |
|  | | | | Selective Logging X Forest | 97.20 | 19.46 | 0.00 |
